# Short-range, orientation-reversing template-switching events occur at a high frequency in the human and yeast genomes

**DOI:** 10.1101/2020.03.06.980458

**Authors:** Samuel Tremblay-Belzile, Juliana Pérez Di Giorgio, Audrey Loubert-Hudon, Alain Verreault, Normand Brisson, James G. Omichinski

## Abstract

The identification of structural variations in genomes using next-generation sequencing approaches greatly facilitates the study of genetic and genomic diseases. The data generated using these approaches also provide interesting new means to examine DNA repair, recombination, and replication to better understand sources of genomic instability. To better utilize this data, we developed SCARR (Systematic Combination of Alignments to Recreate Rearrangements) to identify DNA rearrangements, and used it to examine the occurrence of orientation-reversing events in human and budding yeast genomes. SCARR exceeds the sensitivity of previous genome sequencing approaches, and identifies rearrangements genome-wide with base-pair resolution, which helps provide insights into the mechanisms involved in their formation. We find that short-range orientation-reversing events occur at high rates in both human and yeast genomes. We quantified these rearrangements in yeast strains lacking various DNA repair factors, and propose that these short-range events often occur through template-switching events within a replication fork. We hypothesize that this mechanism may act as an error-prone alternative to fork reversal to restart stalled replication forks.

## INTRODUCTION

Genomic instability results from the inability of DNA processes to perfectly preserve the native genome sequence. This may result in single nucleotide variants when only one base is changed, or structural variants (SVs) when longer sequences are affected. SVs include deletions, duplications, inversions and translocations, all of which have been linked to a variety of genomic diseases [1]. The study of genomic diseases involves the identification not only of SVs [2], but also of the mechanisms that lead to their formation [1]. These include non-allelic homologous recombination (NAHR), which involves long homologous sequences [3], and non-homologous end-joining (NHEJ), which requires no sequence homology [4]. Other mechanisms, such as microhomology-mediated recombination (MHMR) and alternative end-joining (alt-EJ) involve short similar sequences termed microhomologies [5,6]. Although these mechanisms predominantly occur as a result of double-strand breaks (DSBs), genomic instability can also result from replication-dependent mechanisms that do not require breaks in the DNA backbone [7]. Deletions, expansions and duplications may occur as a result of replication slippage, where the DNA polymerase jumps to a different position on the same template strand, whereas fork stalling and template switching (FoSTeS) can lead to more complex rearrangements [7]. These complex events pose additional challenges for detection methods, and are therefore not as well understood as other types of SVs.

Studies have previously reported SVs resulting from FoSTeS during which the nascent leading strand anneals to the lagging strand template near the replication fork. In budding and fission yeast, this annealing step is favored by nearby inverted homologies, and leads to the formation of isochromosomes when replication proceeds uninterrupted after changing direction [8–10]. These complex SVs, termed replication U-turns by Mizuno et al. (2013), were initially identified using reporter systems in nuclear genomes, but they have also been detected in the genomes of plastids and mitochondria using next-generation sequencing [11]. This type of approach revealed that these events in organelles are also favored by sequence similarities as short as 5 bases, and may even occur in the absence of homology. The circumstances leading to their formation, as well as their possible consequences, remain poorly understood due to the challenges involved in their detection by traditional biochemical methods [12].

The rapidly expanding repositories of next-generation sequencing data have opened several new avenues for understanding genomic instability including its roles in evolution and genomic diseases. A landmark study in the field identified tens of thousands of SVs within the human genome [2], which included deletions, duplications, insertions, as well as a small number of inversions. New computational approaches to detect and investigate SVs continue to be developed to improve detection rates and focus on specific types of SVs, such as the harder to detect inversions [13–15]. Nevertheless, complex SVs such as those that result from FoSTeS and other template-switching events often remain undetected by current next-generation sequencing analysis pipelines.

To avoid a bias for simple SVs, we developed SCARR (Systematic Combination of Alignments to Recreate Rearrangements), which uses no *a priori* assumptions about the relative positions of DNA sequences around a breakpoint. SCARR instead iterates through all possible genome-wide alignment combinations to find the best match. Using this approach, it is possible to identify deletions, tandem duplications and breakpoints resulting from template-switching events at base-pair resolution with reasonable accuracy. Consequently, SCARR is particularly well suited for the detection of rare events, such as SVs that result in loss of cell viability. This allows for the study of DNA rearrangements based on a number of parameters, such as microhomology usage, base-pair distance between the rearranged sequences, or SV type. In addition to linking specific SVs to phenotypes, our strategy can also examine how specific factors (DNA repair proteins, replication stresses, DNA damage) impact DNA stability in greater detail. In this study, we used SCARR on datasets from healthy human tissues as well as from several yeast mutant strains exposed to a range of stress conditions to examine patterns of SVs and mechanisms linked to short- and long-range orientation-reversing events.

## RESULTS

### SCARR sensitivity exceeds previous approaches to detect deletions, duplications, and inversions

To determine whether SCARR provides more sensitive detection of DNA rearrangements than previously-published computational methods, we used SVsim and WGSIM to generate simulated next-generation sequencing datasets of a genome containing rearrangements, as previously described [16]. SVsim generates rearrangements in a reference genome, and WGSIM simulates sequencing data from the resulting file. We then identified SVs in the simulated datasets using SCARR and LUMPY to compare their sensitivity and rate of false discovery at low coverage. Even at 1X sequencing coverage, SCARR identifies over 40% of deletions, duplications, and inversions, whereas the false discovery rate remains around 10% (Supplemental Fig S1). In contrast, LUMPY has a much higher sensitivity for inversions (23.7%) than for deletions (4.3%) and duplications (5.2%), but nevertheless only reports less than 25% of total inversions. Since SCARR identifies all rearrangements at base-pair resolution, the data can be used quantitatively. Its actual sensitivity reaches close to 50% when rearrangements that are identified multiple times are taken into consideration. As an additional control, we performed the same analysis on simulated data containing no SVs. These studies revealed that approximately half of the false positives generated by SCARR correspond to genuine SVs being misidentified, whereas the other half arise as a result of mapping ambiguities in the reference genome independently of any SVs (Supplemental Fig S2).

To test the accuracy of SCARR in identifying template switching events, we developed and used SimulateFoSTeS to generate 5X coverage paired-end sequencing in which FoSTeS events occur every 1000 read pairs (See Materials and Methods for details). Unlike SVsim and WGSIM, SimulateFoSTeS does not rearrange the reference genome prior to sequencing, but rather produces unique template-switching events from random positions on the given reference genome at a given rate. Of the 50,188 unique events simulated, SCARR was able to correctly identify 26,846 reads containing SVs, producing no false positives in the process (Supplemental Table S6). This sensitivity of 53.5% is consistent with the results obtained for other types of SVs using SVsim and WGSIM (Supplemental Fig S1). Of the 26,846 SVs identified, 21,349 (79.5%) perfectly matched both fragments of the breakpoint junction, and 5048 (18.8%) perfectly matched one fragment of the breakpoint junction (Supplemental Table S6). Non-matching reads often occurred as a result of repeated sequences in the human genome. This level of accuracy suggests that SCARR can be used to detect patterns of genome instability using rare events at low genome coverage.

To test SCARR on a real dataset, we analyzed public datasets from whole genome sequencing of healthy human brain and liver tissues as paired-end reads of 101 bases. We identified a surprisingly high number of rearrangements for the sequencing coverage of the initial datasets, with over 17,000 rearrangement junctions per genome copy on average (Table 1 and Supplemental Tables 1-2). Following detection, SCARR classifies rearrangements as deletions, duplications, inversions or translocations. Since inversions are detected as single breakpoints for the most part, they include orientation-reversing events, in which two sequences of opposing orientation from the same chromosome are joined, as well as true inversions, in which a sequence is replaced by its reverse complement. For each dataset, we found that over 7000 SVs per genome copy are either deletions or duplications of less than 50 bp, and this accounts for approximately 40% of the SVs detected. Interestingly, the number of each type of rearrangement relative to coverage is very similar between the two datasets, with the exception of inversions, which are approximately 50% more abundant in the brain dataset.

**Table 1.** Summary of coverage and rearrangements for all human datasets.

|  | Brain 1 | Liver | Brain 2 | Spleen |
| --- | --- | --- | --- | --- |
| Read length (bases) | 101 | 101 | 151 | 151 |
| Genome coverage | 13.42 | 12.21 | 6.17 | 5.57 |
| Rearrangements |  |  |  |  |
| Total | 236728 | 201932 | 343899 | 273099 |
| Unique | 170292 | 142117 | 303107 | 235834 |
| Total (per genome) | 17638.9 | 16527.9 | 55741.6 | 49042.0 |
| Unique (per genome) | 12688.7 | 11632.1 | 49129.7 | 42350.1 |
| Microhomology ≥ 5 bp (%) | 84.8 | 85.2 | 55.2 | 68.7 |
| Total del/dup<br>< 50 bp (per genome) | 7038.0 | 7017.5 | 11573.0 | 11715.7 |
| Deletions (per genome) |  |  |  |  |
| Total | 5900.0 | 5789.5 | 9811.9 | 9983.2 |
| Unique | 2996.0 | 2965.0 | 6969.7 | 6907.2 |
| Duplications (per genome) |  |  |  |  |
| Total | 2780.0 | 2792.7 | 6289.0 | 6276.0 |
| Unique | 1893.2 | 1863.4 | 5201.2 | 5024.4 |
| Inversions (per genome) |  |  |  |  |
| Total | 3084.0 | 2056.8 | 5865.3 | 5656.8 |
| Unique | 2866.6 | 1877.5 | 5470.8 | 5271.8 |

Microhomology usage represents the proportion of total rearrangements with at least 5 bases of homology at the breakpoint junction. The number of unique rearrangements is obtained by considering identical events as a single rearrangement. Total del/dup < 50 bp represents total deletions and duplications of sequences shorter than 50 bp.

To determine the effect that read length has on rearrangement detection using SCARR, we sequenced DNA from healthy human brain and spleen tissues as paired-end reads of 151 bases. These datasets yielded approximately twice as many deletions, duplications, and inversions per genome copy as the shorter reads from the publicly available datasets (Table 1 and Supplemental Tables 3-4). This improvement in sensitivity is most likely the result of better sequence alignments in longer reads, which helps compensate for the higher mutation rates found at rearrangement junctions. The added sequence length also increases the chance of successful BLAST+ alignments that do not overlap, with a less pronounced impact on the overlapping reads that result from sequence homology. This leads to a higher proportion of junctions that share little to no homology when longer reads are used (Table 1).

### Short-range orientation-reversing events are frequent in the human nuclear genome

Since SCARR is based in part on an earlier approach that proved very useful in identifying replication U-turns in organelle genomes [11], we also examined the occurrence of these events in the human nuclear genome (Fig 1A). To do this with SCARR, we therefore calculated total orientation-reversing events for each distance between 0 and 200 bases and normalized them to genome coverage (Fig 1B). In all samples, we found that a large majority of short-range orientation-reversing events occur at distances under 50 bases, with a maximum peak at distance 0.

**Figure 1.**
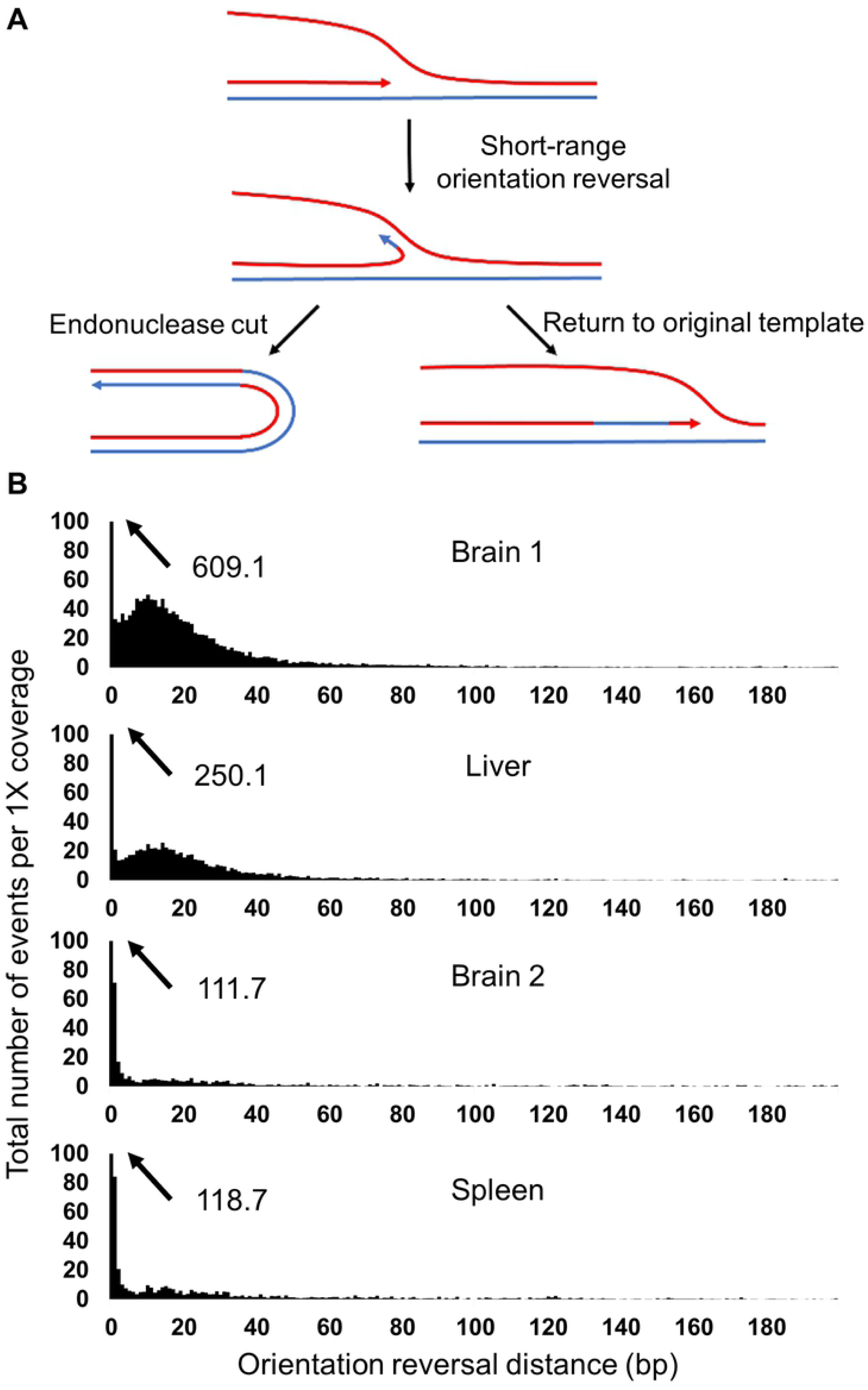
Patterns of short-range orientation-reversing events in human datasets. (**A**) Model for short-range template-switching events within a replication fork leading to orientation reversal. (**B**) Histograms represent the total number of events identified for each distance value between 0 and 200 bp for each dataset, normalized to 1X genome coverage. Datasets are labeled according to the names in Table 1, with read lengths in parentheses for datasets derived from the same DNA samples. Values that exceed 100 are indicated with arrows and their numerical value.

When SCARR fails to explain a read as a combination of two shorter alignments, the script then attempts to explain it as a combination of three alignments. These SVs, which we refer to as paired rearrangements, can provide additional context to rearrangements within a single DNA molecule. Since uninterrupted replication following orientation reversal results in acentric or dicentric chromosomes [9], they can result in large sequence alterations that would be deleterious to the cell. We therefore investigated paired rearrangements in human datasets with 151-bp reads to determine whether or not orientation-reversing events are followed by a second switch that restores replication in the original direction. Since this requires two rearrangements within one read length, the brain and spleen datasets yielded only 920 and 802 total paired rearrangements (Supplemental Tables 3-4), compared to 343,899 and 273,099 single rearrangements, respectively.

Fifteen of the paired rearrangements in the brain dataset and fourteen in the spleen dataset are paired inversions, in which at least one occurred at less than 50 bp (Supplemental Tables 3-4). Interestingly, 5 of the total paired orientation-reversing events were identified 2 to 4 times on independent DNA fragments. Considering the sequencing coverage of approximately 6X for each dataset, these independently-identified paired inversions likely correspond to heterozygous alleles in the individuals from which the samples were obtained. In all cases except one, the longest distance of the two events is less than 400 bp, suggesting that either the nascent strand reanneals to its original template following a template switch or that a second orientation-reversing event occurs. The SVs produced can be complex, creating tandem inverted duplications which result in some DNA segments being triplicated (Fig 2A). In other cases, the SVs are almost perfect true inversions of short sequence fragments (Fig 2B). Given their tendency to form hairpins, we have been unable to validate short-range orientation-reversing events by PCR. However, we were able to amplify and sequence three of the paired orientation-reversing events to confirm the alignments obtained with SCARR, which also supports their presence as allelic variants (Supplemental Fig S3).

**Figure 2.**
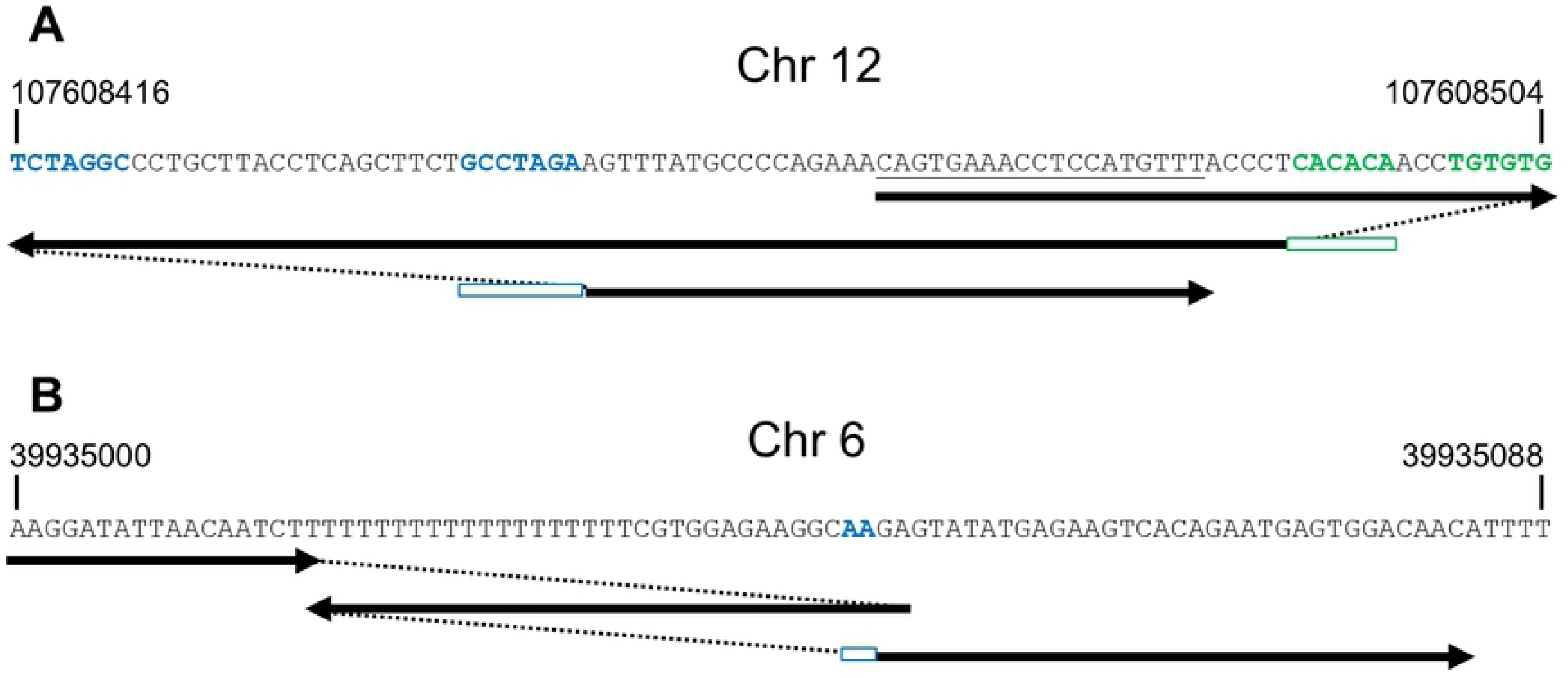
Examples of paired orientation-reversing events occurring at short range identified in the spleen dataset. The arrows follow DNA replication, with dotted lines representing template switches and changes in orientation. The nascent strand adds the sequence presented when replication goes from left to right, and its reverse complement when replication goes from right to left. The product sequence is determined by following the arrows, with jumps along the dotted lines. Inverted microhomologies that may contribute to the template switching are represented in blue and green, with rectangles of corresponding color where the annealing would lead to template switching. The underlined sequence is triplicated as a result of the two inversions.

### Yeast mutant strains as a model to study short-range orientation-reversing events

To explore the proteins and factors involved in short-range orientation-reversing events in eukaryotic nuclear genomes, we sequenced DNA from several strains of *Saccharomyces cerevisiae*, including some with deletions in genes involved in DNA metabolism. To measure the impact of specific repair pathways on orientation reversal events, we used mutants for genes involved in end joining (*dnl4-Δ* and *yku70-Δ*), long-range end resection (*exo1-Δ* and *sgs1-Δ*), end processing (*mre11-Δ* and *sae2-Δ*), homology search and single-strand DNA binding (*rad51-Δ, rfa1-S373P* and *mgs1-Δ*), as well as DNA-damage sensing and replication (*srs2-Δ, mph1-Δ, rad9-Δ* and *pol32-Δ*). We grew each strain in four different conditions to test the effect of replication stress and DNA damage: YPD culture medium, α-factor (αF) followed by camptothecin (CPT), nocodazole (NOC) followed by CPT, and hydroxyurea (HU). CPT forms a ternary complex with topoisomerase I and a nicked DNA duplex, whereas αF and NOC stop cells during either the G1 or M phase, respectively. Lesions caused by CPT form DSBs when they are encountered by a replication fork. CPT treatment on cells arrested with NOC is therefore expected to have little impact on genome stability. HU depletes the nucleotide pool and induces replication stress as well as single-strand breaks [17].

We used SCARR to identify rearrangements in the yeast datasets, and analyzed the patterns for inversions under 1 kb (Fig 3 and Supplemental Table 5). As with the human datasets, we observed that orientation reversal events form a distribution for distances under 50 bases in all datasets, although CPT and HU stresses also showed a larger increase in long-range events. The similarity between patterns for short-range events in these datasets suggests that budding yeast can serve as a model for studying orientation reversal in humans. To identify proteins involved in this pathway, we compared the number of short-range orientation reversal (< 50 bases) and long-range orientation reversal events (≥ 50 bases) in the different yeast strains under the four growth conditions. For each mutant, we calculated the total number of these events normalized to genome coverage for each dataset (Fig 4). These results indicate that short- and long-range events are affected differently by the deletion mutants under the different growth conditions, suggesting that they are mechanistically distinct from each other.

**Figure 3.**
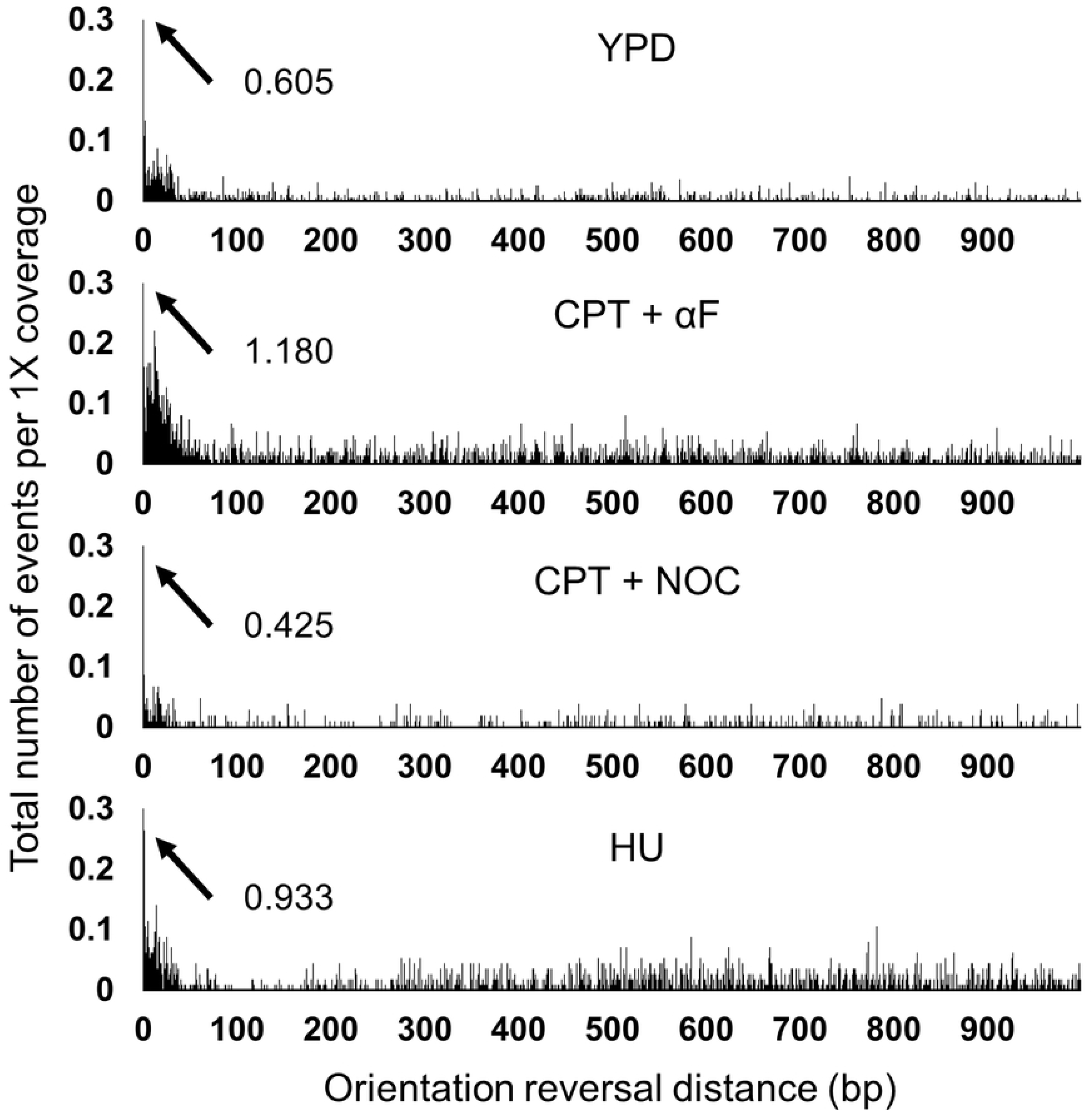
Patterns of short-range orientation-reversing events in the WT yeast strain in the four growth conditions. Histograms represent the total number of inversions identified for each distance value between 0 and 999 bp for each dataset, normalized to 1X genome coverage. YPD medium, hydroxyurea (HU), camptothecin and α-factor (CPT + αF) and camptothecin and nocodazole (CPT + NOC). Values that exceed 0.3 are indicated with arrows and their numerical value.

**Figure 4.**
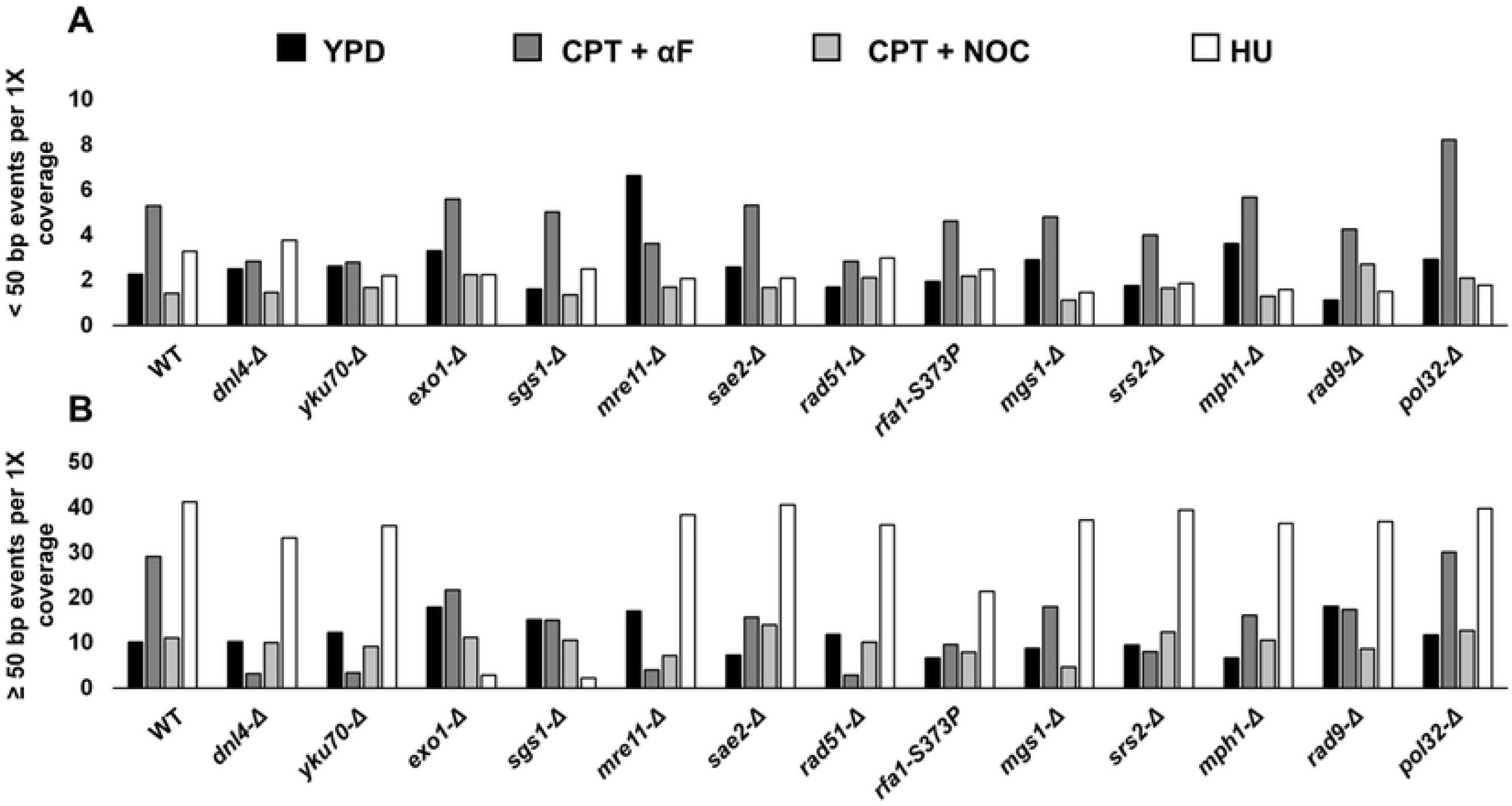
Comparison of short- and long-range orientation-reversing events in the WT and yeast deletion strains under the four different conditions. (**A**) Total number of short-range events, normalized to 1X genome coverage. (**B**) Total number of long-range events, normalized to 1X genome coverage. YPD: YPD medium, αF: α-factor, CPT: camptothecin, HU: hydroxyurea.

### End-joining mechanisms contribute only partly to short-range orientation-reversing events during G1 phase in yeast

Since Dnl4 (DNA ligase IV) and Yku70 (subunit of the DSB-binding Ku complex) are both closely linked to the NHEJ [18], we observed similar results for the *dnl4-Δ* and *yku70-Δ* strains under all four growth conditions (Fig 4). Since NHEJ is mostly active before S phase, we obtained the most striking results for these two strains in the presence of CPT + αF. Although these datasets showed a decrease in short-range orientation reversal compared to the wild-type (WT) strain, they also yielded some of the lowest numbers for long-range orientation reversal. In contrast, all other conditions showed very minor difference between the WT, *dnl4-Δ* and *yku70-Δ* strains, suggesting that HU stress mostly creates DSB ends between S and M phase, when recombination pathways are active. These results suggest that most long-range events and some short-range events require the end-joining machinery during the G1 phase when DSB end-resection is suppressed. However, the milder decrease in short-range orientation-reversing event in these conditions suggests that they also occur through an alternate mechanism that requires neither end-joining nor end-resection.

### Extensive end-resection leads to long-range orientation reversal under HU stress

Since Exo1 and Sgs1 are involved in extensive DSB end resection, which is necessary for homologous recombination (HR) [19], their absence was expected to lead to an increase in rearrangements from error-prone mechanisms after NOC treatment. Indeed, the *exo1-Δ* and *sgs1-Δ* strains both showed increases in long-range orientation reversal in the absence of stress. However, these strains presented almost no difference from the WT strain under CPT + NOC stress, with only a modest increase in short-range orientation reversal in the absence of Exo1 (Fig 4). The increase in long-range events observed in the absence of stress therefore seems to occur outside of the G2/M phase. Interestingly, the *exo1-Δ* and *sgs1-Δ* strains also showed a dramatic decrease in long-range events under HU stress (Fig 4 and Supplemental Fig S4). The distance distribution for the Exo1- or Sgs1-dependent events observed in the WT strain falls well within the expected resection range of the two nucleases [19]. This is consistent with an increase in snap-back DNA synthesis following long resection involving Exo1 or Sgs1 under HU stress.

### The impact of Mre11 on short-range orientation reversal events seems independent of its role in end resection

The Mre11-Rad50-Xrs2 (MRX) complex and Sae2 are known to participate in the initiation of resection by Exo1 [20–22]. As such, we expected the *mre11-Δ* and *sae2-Δ* strains to show similar effects as the *exo1-Δ* strain. However, the *sae2-Δ* strain yielded similar levels of orientation reversal to the WT strain under all four conditions, except for a reduction in long-range events with CPT + αF stress, similar to the *exo1-Δ and sgs1-Δ* strains (Fig 4). In contrast, the *mre11-Δ* strain presented short-range orientation reversal levels that are noticeably different from all other mutants (Fig 4). The *mre11-Δ* strain displayed more short-range events than any other yeast deletion strain in the absence of stress, but fewer short-range events than the WT strain under HU and CPT + αF stresses. The *mre11-Δ* strain also presents a reduction in long-range events comparable to the *dnl4-Δ* and *yku70-Δ* strains in the presence of CPT + αF. These differences between *mre11-Δ* and other mutants involved in end resection suggests Mre11 might play a distinct role in short-range orientation reversal events.

### Short-range orientation-reversing events do not seem to occur through recombination

In a previous study, we found that the single-stranded DNA-binding Whirly proteins and the bacterial-type RecA recombinase played important roles in preventing replication U-turns in plastids of *Arabidopsis thaliana* [11]. Rfa1 (subunit 1 of the replication protein A complex) and Rad51 both bind single-stranded DNA as part of the HR pathway [23], with Rfa1 also inhibiting microhomology-mediated rearrangements [24]. The deletion of *RFA1* is lethal in yeast, but the *rfa1-S373P* mutation (also known as *rfa1-t33*) increases genome instability while maintaining cell viability [25]. In our datasets, the *rad51-Δ* and *rfa1-S373P* strains showed some similarities, including modest increases in short-range inversions but no increases in long-range inversions, when compared to the WT under CPT + NOC stress (Fig 4). Under CPT + αF stress, the *rad51-Δ* strain presented the same pattern as the *dnl4-Δ, yku70-Δ* and *mre11-Δ* strains, which would support previously-reported roles in end-processing for Mre11 [26] and Rad51 [27].

Mgs1 possesses single-stand annealing activity, and the *mgs1-Δ* mutation was shown to cause an increase in recombination and genome instability after replication [28]. Surprisingly, our datasets with the *mgs1-Δ* strain showed a decrease in long-range orientation reversal compared to the WT strain under CPT + NOC and CPT + αF stresses, with little change to short-range events (Fig 4). The opposite was observed in the presence of HU, with long-range events maintaining a level similar to the WT, but short-range events being reduced by more than half. These effects are consistent with different roles for Mgs1 in the presence or absence of DNA-damage [29].

The helicases Srs2 and Mph1 both favor the synthesis-dependent strand-annealing (SDSA) pathway of homologous recombination [30]. Srs2 also inhibits recombination by dismantling Rad51 filaments. In our results, neither the *srs2-Δ* nor the *mph1-Δ* strain showed any large variation from the WT strain under CPT + NOC stress, suggesting a limited impact for homologous recombination pathways on orientation reversal events.

### Replication stress contributes to short-range orientation reversal events

The DNA-damage checkpoint protein Rad9 prevents DNA-degradation at stalled replication forks [31]. In the absence of stress, the *rad9-Δ* strain presented the lowest level of short-range inversions, but the highest level of long-range events (Fig 4). Compared to the WT strain, this strain displayed fewer short-range orientation reversal events under HU stress, but more under CPT + NOC. The effect of Rad9 on short-range events therefore seems to be different between conditions of replication stress and DNA damage.

Pol32 is an error-prone polymerase that was found to be responsible for replication during repair of broken forks [32]. In the four growth conditions, the *pol32-Δ* strain presented similar levels of long-range orientation reversal events as the WT strain. However, it showed more short-range events than the WT strain in both CPT treatments, and fewer in the presence of HU. Pol32 therefore appears to prevent rearrangements in the presence of DNA damage, while promoting instability during replication stress. This would be consistent with Pol32 activity favoring replication following template-switching events.

### Short-range orientation reversal events require little to no sequence homology

Replication U-turns were shown in previous studies to occur at sites of inverted repeats through the annealing of short homologous sequences [9,10]. The data obtained from SCARR also includes patterns of sequence similarity at rearrangement junctions, which can help distinguish between different rearrangement mechanisms and provide information about their requirements for sequence similarity. To determine whether orientation reversal requires inverted repeats near the breakpoint, we investigated the length of sequence similarity involved in short-range events in our data. Since we are mostly interested in short-range events that occur through mechanisms that do not produce long-range events, we looked in more detail at yeast deletion mutants and stress conditions that suppress long-range orientation reversal. These conditions are also likely to suppress short-range events occurring through the same mechanisms, and should therefore yield a pattern of sequence similarity requirement more specific to short-range template-switching events. We then compared these mutants to the WT strain without induced stress.

In the WT dataset, we observed two main peaks with similar profiles for both short- and long-range orientation reversal events: at 0 bases of homology, and between 4 and 15 bases of homology (Fig 5A). In each case, the two peaks reach approximately the same maximum values. In contrast, the *dnl4-Δ, yku70-Δ, mre11-Δ* and *rad51-Δ* strains in the presence of the CPT + αF stress all yielded the same peaks, but with a higher relative proportion for short-range events with 0 bases of homology compared to homologies between 4 and 15 bases (Fig 5B and Supplemental Fig S5). The same pattern is also observed for the *exo1-Δ* and *sgs1-Δ* strains under HU stress (Fig 5C and Supplemental Fig S5). Interestingly, the human datasets with reads of the same length also presented the same pattern for short-range events, in spite of having noticeably different patterns for long-range events (Fig 5D). These results further support a distinct mechanism for the short-range orientation-reversing events that occur in the human nuclear genome, and suggest that sequence homology is not a requirement in this mechanism.

**Figure 5.**
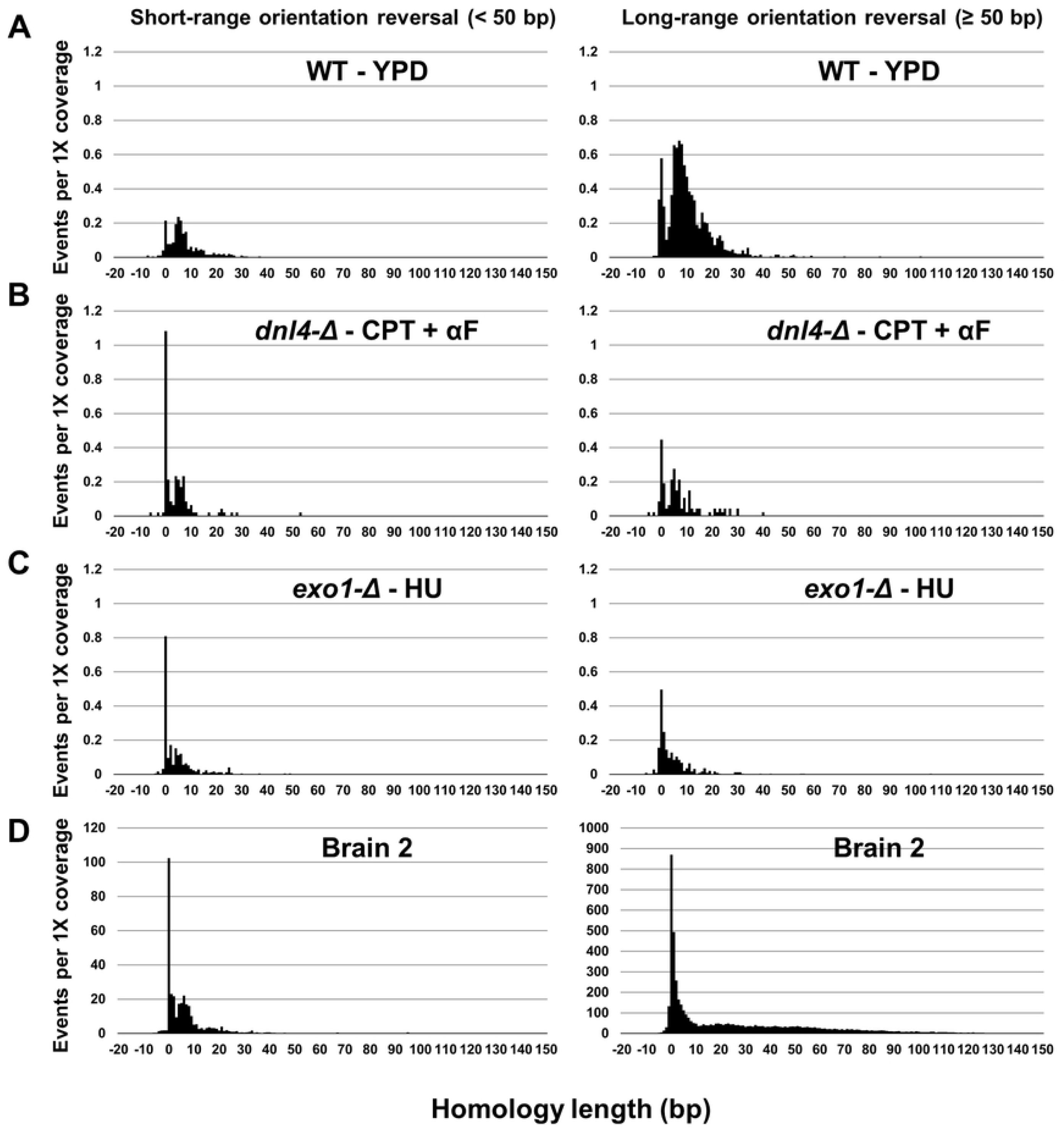
Homology usage for short-range (< 50 bp) and long-range (≥ 50 bp) orientation-reversing events. Histograms represent the total number of unique events for which a homology of given length is found at the breakpoint junction. Negative homology lengths represent base insertions at the breakpoint junction. All values are normalized to 1X genome coverage. (**A**) WT yeast strain grown in YPD. (**B**) *dnl4-Δ* strain grown in CPT (5 μg/mL) and αF. (**C**) *exo1-Δ* strain grown in HU (200 mM). (**D**) Human brain 2 dataset.

## DISCUSSION

Replication U-turns were initially observed in fission yeast using a reporter system with perfect 2.6 kb inverted repeats, and were detected with repeats as short as 150 bp [9,10]. From these results, it was suggested that shorter homologies may be sufficient for U-turns to occur at an appreciable rate. By analyzing the genome-wide occurrence of short-range orientation reversal events in budding yeast, we observe that a significant proportion of these rearrangements occur independently from end-joining mechanisms and in the absence of end resection. This is consistent with a replication-based template-switching mechanism. We also note that these short-range orientation-reversing template-switching events may be favored when the nascent strand anneals to the new template using inverted repeats, but that they also occur frequently in the absence of sequence homology at the breakpoint junction. These results support a model where replication U-turns form when a template switch occurs before a stalled fork collapses. The availability and proximity of single-stranded DNA within a replication fork could explain their frequent occurrence in the absence of sequence similarity. It is possible that the nascent strand uses a stretch of sequence similarity a few bases away from its extremity to anneal to a new template and resume replication using a low-fidelity polymerase.

We also report the detection of short-range orientation-reversing template-switching events in the human nuclear genome with distance and homology usage patterns that closely resemble those observed in yeast. In some cases, two such orientation reversal events are paired to return the DNA strand to its original orientation. A similar mechanism was recently proposed to explain mutation clusters observed between chimpanzee and human reference genomes, as well as between the assembled genomes of individual humans [33]. The rate at which we identified short-range orientation reversal events in healthy human tissue further supports the idea that they play a role in both the evolution of genomes and the appearance of genetic diversity within species. Interestingly, similar rearrangement patterns as described in Fig 2A, but spanning several kb or Mb, have been reported by other groups and termed duplication-inverted triplication-duplication [34–36]. These events were proposed to occur via a recombination event followed either by an end-joining or break-induced replication event. However, considering the large difference in scale between these rearrangements and those identified by SCARR, it is unclear whether they arise from the same mechanism.

The presence of U-turns in the human nuclear genome also suggests a new mechanism by which stalled replication forks can be restarted (Fig 6). A previous study has found that the hypersensitivity of BRCA2-deficient cells to DNA-damaging agents depends on the recruitment of MRE11 to stalled replication forks [37]. BRCA2, together with RAD51 and the SMARCAL1 helicase, participates in the conservative restart of stalled forks through fork reversal [38]. Though replication U-turns result in alterations in the genome sequence, they also allow DNA synthesis to resume without creating DSB intermediates. Combined with our results in yeast, this raises the possibility that this type of mechanism may play a role for the chemoresistance observed in BRCA2-deficient cells. Since HR is impaired in BRCA2-deficient cells, fork restart through U-turns could explain a reduced sensitivity to fork stalling in the absence of fork reversal. The rate at which we observe short-range orientation-reversing template-switching events in healthy tissues and the proposed evolutionary model also suggest that the SVs resulting from U-turns can occur with little to no deleterious effects [33].

**Figure 6.**
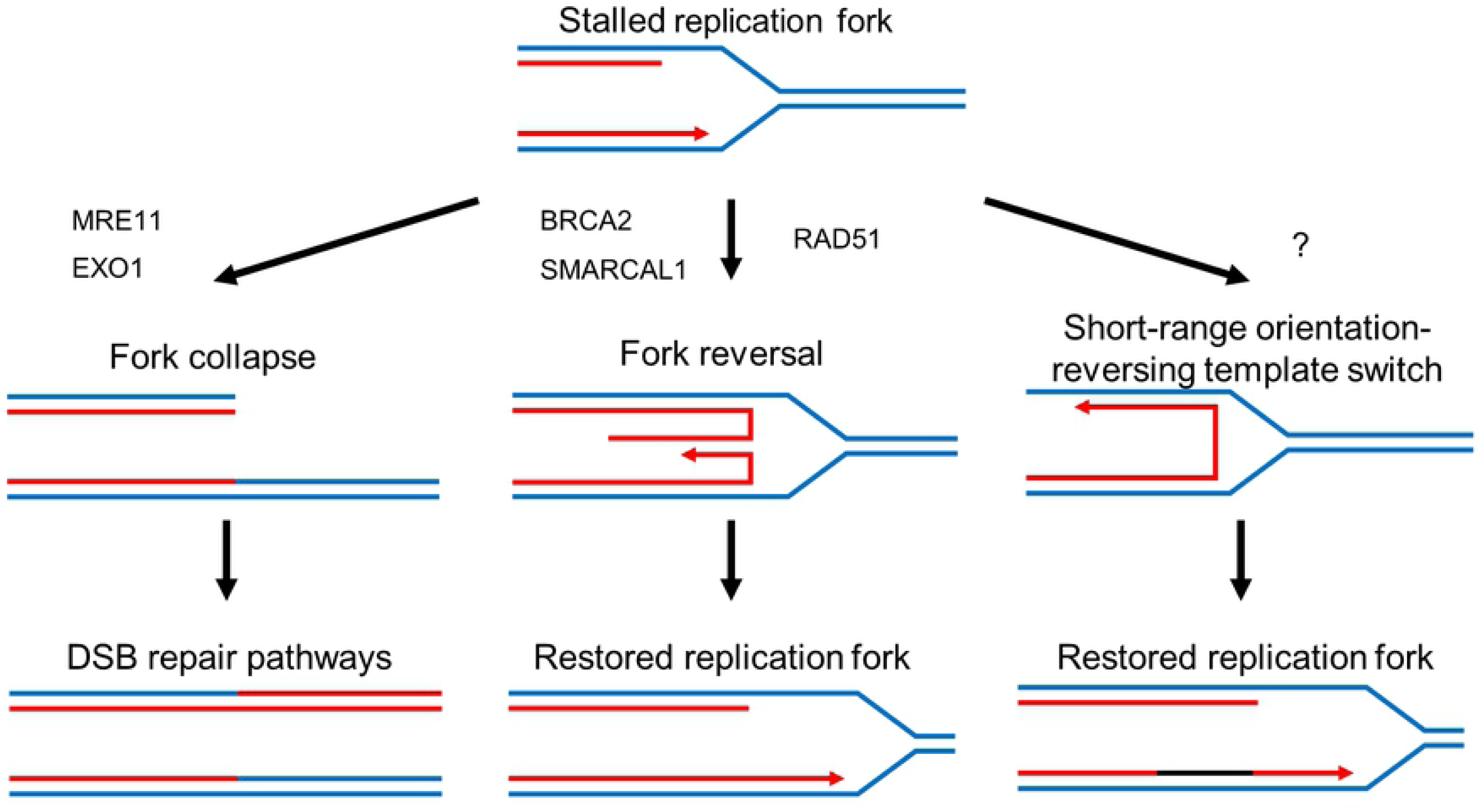
Short-range orientation-reversing template switch as an alternate mechanism to restart stalled replication forks. Blue lines represent parent DNA strands and red lines represent newly synthesized strands. Stalled forks may collapse following nuclease activity and result in double-strand breaks, which are taken over by the DSB repair machinery. Fork reversal may be initiated by SMARCAL1, and involves annealing of the nascent strands by BRCA2 and RAD51. This structure is stabilized by proteins until the lesion causing fork stalling is removed, and the replication fork is restored. In the absence of machinery necessary for fork reversal, the nascent leading strand may switch templates within the replication fork and continue replication on the lagging strand template. After the lesion causing fork stalling is removed, the replication fork can be restored with a mismatched inverted segment of DNA, represented by the black line.

Since datasets with longer reads improve the sensitivity of SCARR, next-generation sequencing technologies that produce longer read lengths will be an important avenue to explore in the future. The current version of SCARR checks each read for either one or two rearrangements, but the approach can be extended to any number of rearrangements per read, provided the sequences are long enough to successfully align a sufficient number of fragments. As such, long reads will also help to provide context for rearrangements within single molecules. This possibility will be particularly interesting in the study of orientation-reversing template-switching events, since they require a second template-switching event to occur in order to resume DNA synthesis in the original direction.

## MATERIALS AND METHODS

### Simulated datasets

SVs were randomly created using SVsim (https://github.com/GregoryFaust/SVsim) to generate 2,500 deletions, duplications, and inversions in the human reference genome (build 38) from which the mitochondrial genome has been removed. The variants each range from 100 bp to 10,000 bp, as described previously [16]. Virtual sequencing was performed using WGSIM (https://github.com/lh3/wgsim) to a coverage of 1X, using paired-end reads of 150 bp and default settings for insert size and error rates, also as per Layer et al. 2014. Rearrangements with lengths that differ by more than 20 bases from the simulated SV lengths were classified as false positives. The same sequencing was also performed on the unmodified human reference genome GRCh38 (minus the mitochondrial genome), with default error and indel settings. Rearrangements were detected with SCARR 1.0 and LUMPY 0.2.13.

### Public datasets

Datasets for healthy human brain (ERR419275, ERR419276, ERR419277, ERR419278) and liver (ERR419279, ERR419280, ERR419281, ERR419282) DNA were obtained from the NCBI SRA. Additional library and sequencing information relating to the datasets was previously described [39].

### Human DNA samples

Human DNA was purchased commercially from BioChain, pre-extracted from the following healthy tissues: brain (Occipital lobe, catalog no. D1234062, lot B806171, male, 41 years old) and spleen (Catalog no. D1234246, lot A712149, male, 27 years old).

### Yeast strains

Haploid deletion mutants for *DNL4, EXO1, MGS1, MPH1, MRE11, POL32, RAD9, RAD51, SAE2, SGS1, SRS2* and *YKU70* were obtained from the yeast Magic Marker strains from Open Biosystems, as per their protocol [40]. Successful selection of the deletion mutant was confirmed by PCR using the suggested primers. To obtain *rfa1-S373P* in the same background strain, the point mutation was generated in the *RFA1* plasmid from the Molecular Barcoded Yeast ORF library from Thermo Fisher Open Biosystems, and transformed into the heterozygous *rfa1-Δ* strain from the Magic Marker collection. The haploid deletion mutant containing the plasmid was then obtained by following the protocol for Magic Marker strains but subtracting uracil from the medium.

### Growth conditions and DNA extraction

Cells were grown in YPD to an OD_600_ of 0.5 before treatment. For the CPT + αF treatment, the cells were incubated for 1 h in 5 μg/mL αF (Bachem) to stop the cell cycle. Cell cycle arrest in G1 was confirmed at 97% (863/890) by microscopy. An additional 5 μg/mL αF was then added at the same time as 5 μg/mL CPT (Sigma-Aldrich). The cells were pelleted for DNA extraction 2.5 h after adding the CPT. For the CPT + NOC treatment, the cells were incubated for 2 h in 15 μg/mL NOC (Sigma-Aldrich) before adding 5 μg/mL CPT. Cell cycle arrest in M phase was confirmed at 95.5% (1266/1325) by microscopy. The cells were pelleted for DNA extraction 2.5 h after adding the CPT. For the HU treatment, 200 mM HU (BioShop Canada) was added, and the cells were pelleted for DNA extraction after 3 h. For the control, 1µl/mL dimethyl sulfoxide (DMSO) was added, and the cells were pelleted for DNA extraction after 3h. DNA was extracted by phenol/chloroform and treated with RNAse A.

### Illumina sequencing

Libraries were prepared from the DNA samples using the NxSeq AmpFREE Low DNA Library kit (Lucigen, Cat no. 14000-2) as per the manufacturer’s protocol. This includes SPRI bead cleanup and size selection steps for a final median insert size of approximately 320 bp. Sequencing was performed on the Illumina HiSeq X Ten (302 cycles, paired-end). Library preparation and sequencing were done at Génome Québec (McGill University, Montréal, Qc, Canada). All new datasets were made available on NCBI SRA (Project numbers: SRP134058 and SRP154401).

### Confirmation of SVs by Sanger sequencing

Two SVs were amplified from the spleen DNA sample by PCR (SV1 forward primer: ATCGTAATAAATTTTCAGAAGTCCGTGAAA, reverse primer: CTCTCTAAGCCTCAATGTCCTCAG; SV2 forward primer: AAATTAGCCCGCAGGCGTG, reverse primer: CCTGCCCACAGCAATGTGA). The bands corresponding to SVs were gel-extracted and TOPO cloned for Sanger sequencing.

### Rearrangement detection

Datasets were enriched for rearrangement-containing reads using a Galaxy workflow adapted from a previously published approach [11]. Adapter sequences were removed using Trim Galore! (https://github.com/FelixKrueger/TrimGalore) prior to alignment. The sequences obtained were further filtered to remove potential artifacts generated during library preparation from single-stranded DNA [41]. Reads were removed when the first 12 bases of paired reads were identical with 1 mismatch or less and a maximum offset of 3 bases. The remaining reads were labeled as potential junctions and aligned with BLAST+ [42] using the following command: “blastn -query potential_junctions.fasta -db reference_genome.fasta -out output.txt -word_size 10 -evalue 0.0001 -outfmt 6”. To keep file sizes manageable and accelerate the analysis, potential junctions were split into files containing 25 sequences or less and the work was parallelized. The output from BLAST+ was given as input to SCARR with default parameters. SCARR is a custom Python script available on GitHub that tests all possible combinations of BLAST+ alignments to determine whether a satisfactory match can be found (Supplemental Fig S6). Additional custom scripts were used to sort and organize the output rearrangement files from SCARR, and are also available on GitHub.

### Rearrangement analysis

For each rearrangement, microhomology length is determined by the number of bases from the original read that are aligned to both sides of the breakpoint. When some bases at the breakpoint fail to align to either side, they are counted as inserted bases. When the alignments map to different chromosomes, the rearrangement is labeled as a translocation, and no further analysis is performed. When both alignments map to the same chromosome, their relative directions are verified. When they are in opposing directions, the rearrangement is labeled as an inversion, and the distance represents the number of bases between the reference genome positions of the last base in the first alignment and the first base in the second alignment, minus any microhomology length. When both alignments are in the same direction, their relative positions are verified. When the breakpoint represents a jump forward, the rearrangement is labeled as a deletion, whereas a jump backwards is labeled as a duplication. When no microhomology is present, the distance is measured from the last base in the first alignment to the first base in the second alignment. When a microhomology is present, its length is subtracted from the distance, so that distance represents the exact number of bases deleted or duplicated in the reference genome.

### SCARR algorithm

SCARR loads all blastn alignments from a read into an array, and determines the minimum and maximum mapped positions of the read. The script then looks for any single alignment that spans most of the read, tolerating a length difference of at most 5 bases. When any such alignment is found, the read is considered not to contain a rearrangement, and the script continues to the next. When no single alignment spans the read, SCARR tests all possible combinations of 2 alignments. For each combination, it determines what percentage of the read between the minimum and maximum mapped positions are covered by the alignments, and subtracts any gaps or mismatches in the individual alignments (weighted by default at 1.25 per base mismatch and 1.5 per base gap). This determines a score out of 100 for the alignment pair. The best score is kept and compared to a set threshold (by default 80), and the best alignment pair is considered as a rearrangement when it exceeds the threshold. For reads where no alignment pair passes the threshold, the process is repeated by testing all possible combinations of 3 alignments. When the best-scoring alignment triplet exceeds the threshold, it is considered as a paired rearrangement and written to a file separate from single rearrangements. When no alignment triplet passes the threshold, the read is considered as unlabeled, and the script moves on to the next.

### SimulateFoSTeS script and comparison to SCARR results

SimulateFoSTeS simulates rare template-switching events from an unmodified reference genome. Simulated template-switching events do not take into consideration any sequence similarity at the breakpoint. Read length, insert size, frequency of template-switching events and number of reads to generate are all user-specified. The script also generates reference files for the positions of every read, and a separate file for the positions of each template-switching event. The dataset generated in this study used default values (5 million reads, 150 bp sequences, 200 to 300 bp inserts, 1 template-switching event per 1000 read pairs), and was run on the GRCh38 reference genome from which the mitochondrial genome was removed. It is important to note that although blastn alignments used by SCARR overlap when sequence similarities are present at the breakpoint junction, SimulateFoSTeS records all read positions in its reference file as if no sequence similarity was present. As such, SCARR alignments are considered to match when the start position of the first alignment and end position of the second alignment match the SimulateFoSTeS reference file.

### Data access

Sequencing datasets have been deposited at the NCBI Sequence Read Archive (SRA) under accession numbers PRJNA437181 and PRJNA481831.

SCARR and associated scripts are available in the GitHub repository (https://github.com/SamTremblay/SCARR). SimulateFoSTeS is available in the GitHub repository (https://github.com/SamTremblay/SimulateFoSTeS).

## ACKNOWLEDGEMENTS

We thank G. Arseneault for her assistance preparing the yeast strains. The raw sequencing data was treated using Galaxy on the public server (usegalaxy.org) and on the Genetics and Genomics Analysis Platform (GenAP). Computations were made on the supercomputer Briarée, managed by Calcul Québec and Compute Canada. The operation of this supercomputer is funded by the Canada Foundation for Innovation (CFI), the ministère de l’Économie, de la science et de l’innovation du Québec (MESI) and the Fonds de recherche du Québec - Nature et technologies (FRQ-NT). This work was supported by scholarships from the National Sciences and Engineering Research Council of Canada (NSERC) and FRQ-NT to S.T.B.

## SUPPLEMENTAL MATERIAL LEGENDS

**Supplementary Figure S1**. Comparison of sensitivity and false detection rate of SCARR and LUMPY based on 1X coverage of simulated data. Sensitivity for unique rearrangements represents the ratio of unique events to the total number of different expected SVs. For SCARR, each simulated deletion and duplication correspond to one expected rearrangement, and each inversion corresponds to two expected rearrangements. Total sensitivity represents the ratio of total events detected to the total number of events expected at this coverage. False discovery rate corresponds to the ratio of total false positives to the total number of events detected.

**Supplementary Figure S2**. False discovery rate in simulated datasets with and without SVs. Values represent the total number of each type of false positive found in 1X simulated sequencing coverage. The SVsim dataset contains deletions, duplications, and inversions, 2,500 of each type. The control dataset uses the unmodified reference genome.

**Supplementary Figure S3**. Confirmation of paired orientation-reversing events by PCR cloning and Sanger sequencing. For each event, the top sequence corresponds to the reference genome, and the bottom sequence corresponds to the SVs identified by SCARR and confirmed by Sanger sequencing.

**Supplementary Figure S4**. Short- and long-range orientation reversal events in the *exo1-Δ* and *sgs1-Δ* yeast strains in YPD and hydroxyurea (HU) compared to the wild-type (WT) yeast strain under the same conditions. Histograms represent the number of events per distance ranges of 100 bp. Labels represent lowest value of the range.

**Supplementary Figure S5**. Homology usage for short-range (< 50 bp) and long-range (≥ 50 bp) orientation reversal events. Histograms represent total number of unique events for which a homology of given length is found at the breakpoint junction. Negative homology lengths represent base insertions at the breakpoint junction. All values are normalized to 1X genome coverage.

**Supplementary Figure S6**. Summary of the approach used to identify rearrangements. (1) Next-generation sequencing data is fed through a Galaxy workflow to obtain DNA fragments for which at least one extremity is not perfectly mapped to the reference genome, as previously described in Zampini et al. 2015. (2) blastn is used to obtain partial alignments for the remaining reads, and (3) the reads that closely match with a single alignment are discarded. (4) For the remaining reads, a rearrangement score is assigned to each possible combination of 2 partial alignments, taking into account the proportion of the read covered and the number of gaps and mismatches in the alignments. (5) The best scoring combination is called as a rearrangement only when it exceeds the selected score threshold.

**Supplemental Table 1**. Full table of SVs detected in the publicly available datasets from a healthy human brain.

**Supplemental Table 2**. Full table of SVs detected in the publicly available datasets from a healthy human liver.

**Supplemental Table 3**. Full table of SVs detected in the sequenced datasets from a healthy human brain.

**Supplemental Table 4**. Full table of SVs detected in the sequenced datasets from a healthy human spleen.

**Supplemental Table 5**. Statistics of all the SVs detected in the datasets from all yeast strains under all four conditions.

**Supplemental Table 6**. Full table of the SVs simulated using SimulateFoSTeS, detected using SCARR, and a comparison of the experimental alignments to the reference alignments.

